# An efficient platform for astrocyte differentiation from human induced pluripotent stem cells

**DOI:** 10.1101/133496

**Authors:** TCW Julia, Minghui Wang, Anna A. Pimenova, Kathryn R. Bowles, Brigham J. Hartley, Emre Lacin, Saima Machlovi, Rawan Abdelaal, Celeste M. Karch, Hemali Phetnani, Paul A. Slesinger, Bin Zhang, Alison M. Goate, Kristen J. Brennand

**Author notes:** These authors contributed equally (Minghui Wang, Anna A. Pimenova, Kathryn R. Bowles, Brigham J. Hartley, Emre Lacin).

## Abstract

Growing evidence implicates the importance of glia, particularly astrocytes, in neurological and psychiatric diseases. Here, we describe a rapid and robust method for the differentiation of highly pure populations of replicative astrocytes from human induced pluripotent stem cells (hiPSCs), via a neural progenitor cell (NPC) intermediate. Using this method, we generated hiPSC-derived astrocyte populations (hiPSC-astrocytes) from 42 NPC lines (derived from 30 individuals) with an average of ∼90% S100β-positive cells. Transcriptomic analysis demonstrated that the hiPSC-astrocytes are highly similar to primary human fetal astrocytes and characteristic of a non-reactive state. hiPSC-astrocytes respond to inflammatory stimulants, display phagocytic capacity and enhance microglial phagocytosis. hiPSC-astrocytes also possess spontaneous calcium transient activity. Our novel protocol is a reproducible, straightforward (single media) and rapid (<30 days) method to generate homogenous populations of hiPSC-astrocytes that can be used for neuron-astrocyte and microglia-astrocyte co-cultures for the study of neuropsychiatric disorders.

**ABBREVIATIONS:**

hiPSC: human induced pluripotent stem cell
NPC: neural progenitor cell

## INTRODUCTION

Astrocytes are the most abundant cell type in the central nervous system, rivaling the diversity of neurons in cellular morphologies, gene expression profiles, developmental origins, physiological properties, functions, and responses to injury and disease^1^. Within the human brain, astrocytes have a variety of essential functions including glutamate biology, axonal guidance, trophic support, inflammatory response and wound healing, formation of the blood brain barrier, and neuronal synapse formation and plasticity^2-4^. Although the full contribution of astrocytes to neurological disease remains unresolved, astrocyte cell-autonomous deficits have been implicated in a variety of neurological disorders^5,6^. The most significant genetic risk factor for Alzheimer’s disease, Apolipoprotein E4 (*APOE4*), is predominantly synthesized and secreted by astrocytes^7^. Furthermore, astrocytes derived from hiPSC- or mouse-based models of amyotrophic lateral sclerosis ^8,910^, Rett syndrome^11^, and Huntington disease^12^ damage neurons in co-culture or after transplantation.

Evolutionarily, the astrocyte-to-neuron ratio increases from low vertebrates to rodents and to primates^13^. Human cortical astrocytes are larger, structurally more complex and diverse, and propagate calcium waves four-fold faster than their rodent counterparts^14^. Transplantation of human glia into mice enhances activity-dependent plasticity and learning^15^. Given the unique biology of human astrocytes, it is critical that improved human-specific cell-based systems be established, to enable the study of human astrocyte function in health and disease.

Because of their ability to model all of the (known and unknown) genetic risk factors underlying neuropsychiatric disease, hiPSCs are routinely used as a source of various types of neurons and astrocytes for study^16^. Current hiPSC-based methods for the differentiation of astrocytes typically rely on either an NPC^17-22^ or oligodendrocyte progenitor cell^23^ intermediate. While it has been widely demonstrated that hiPSCs can be differentiated to functional astrocytes for cell-based models of neuropsychiatric disorders *in vitro*^19-22^ or engraftment *in vivo*^17,18,23,24^, existing methods are slow (up to six months)^17,22,23^ and/or require sorting to reduce heterogeneity^25,26^. Here, we screened a number of published protocols, along with commercially available media for primary human astrocyte culture, identifying a robust and straightforward differentiation protocol for generating highly pure astrocytes from hiPSCs. By co-culture with microglia, we compared the function of primary human fetal astrocytes and hiPSC-astrocytes in assays for neuroinflammatory response, phagocytosis, and spontaneous calcium activity, concluding that hiPSC-astrocytes are highly similar to their primary counterparts. Altogether, our rapid differentiation protocol, co-culture strategy and scalable phenotypic assays will serve as a robust platform for queries of healthy and diseased human astrocytes.

## RESULTS

### 30-day exposure of hiPSC-derived NPCs to commercial astrocyte media is sufficient to robustly generate hiPSC-astrocytes

We first screened 11 different media conditions on forebrain-patterned NPCs^27,28^ derived from hiPSCs (**Table 1**). The screening conditions included different combinations of fibroblast growth factor 2 (FGF2)^20^, ciliary neurotrophic factor (CNTF)^17,22^, bone morphogenetic protein 4 (BMP4)^22,23,29^, fibroblast bovine serum (FBS)^22,29^, neuregulin^22,30^, insulin^31^ and ascorbic acid (AA)^32^, as well as three commercial astrocyte media (ScienCell, Gibco, and Lonza) for the culture of primary human fetal astrocytes (**Table 1**). Screening criteria included immunoreactivity for two classical markers of astrocyte identity, S100β and GFAP^33^, astrocyte morphology, survival, and cell line variability (**Supplementary Table 1; Supplementary Fig.1a**). While other conditions resulted in cell line variation, limited cell proliferation, and expression of neuronal markers (**Supplementary Table 1**), two commercial media, ScienCell and Lonza, yielded the most S100β- and GFAP-positive astrocyte-like cells (**Supplementary Fig.1b-d**). These results were confirmed across four representative NPC lines both by flow cytometry and immunocytochemistry by 30 days (**Fig. 1a**; **Supplementary Fig.1e-g**). Culture of NPCs in both media, when combined with low initial seeding density (nearly single cells: 15,000 cells/cm^2^) and minimal serum exposure (1-2%), resulted in astrocyte morphology within ten days (**Supplementary Fig.1h**); star-shaped astrocyte morphologies were evident within 30 days (**Supplementary Fig.1i**). Although ScienCell and Lonza astrocyte media showed equivalent efficiencies (**Supplementary Fig.1b-d**), ScienCell medium was selected owing to its lower cost and relative simplicity.

**Table 1.**
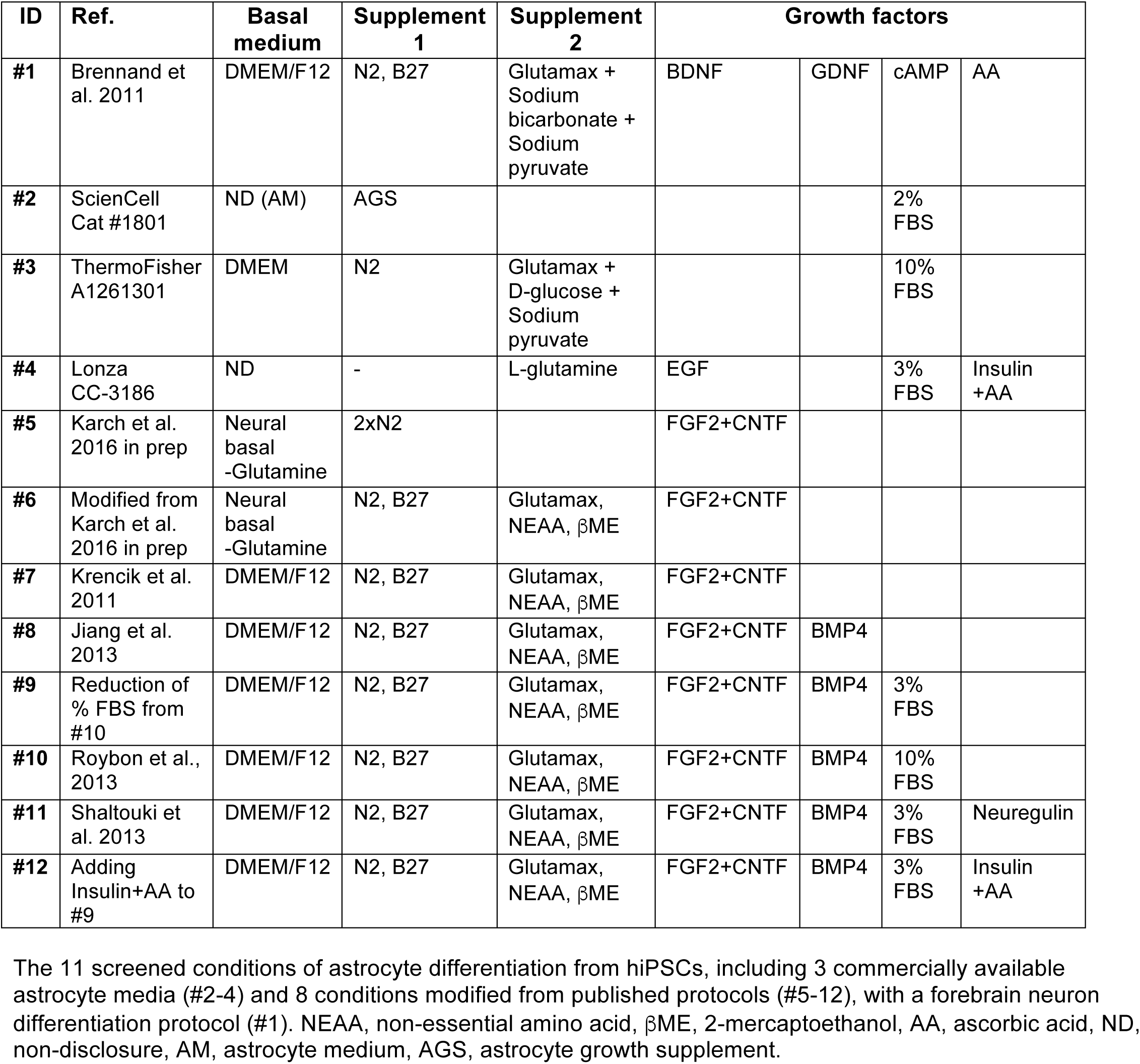
Screening conditions for astrocyte differentiation.

**Figure 1.**
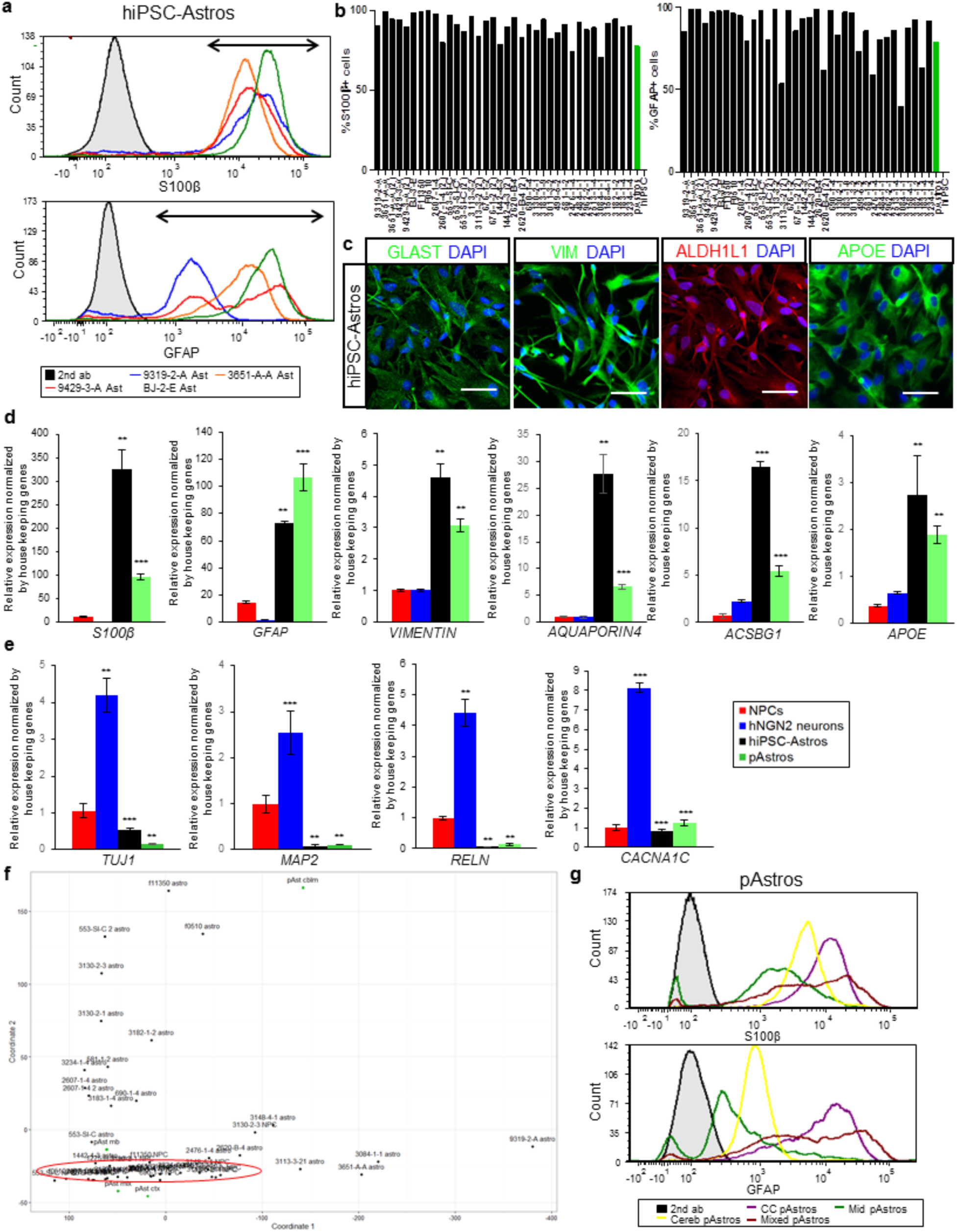
Rapid differentiation of hiPSC-derived NPCs to astrocyte-like identity. (**a**) Representative flow cytometry analysis of S100β (top) and GFAP (bottom) for four 30-day hiPSC-astrocyte differentiations. Arrows indicate the cells positive for each marker protein. (**b**) Graphs of flow cytometry analysis across 35 hiPSC-astrocyte differentiations from 26 NPC lines from three independent hiPSC cohorts. S100β (left) and GFAP (right) immunostaining is shown, with primary human fetal astrocytes (positive control) and hiPSCs (negative control). (**c**) Representative immunofluorescence images of hiPSC-astrocytes stained for astrocyte markers glutamate transporters GLAST (green), VIMENTIN (green), ALDH1L1 (red), and APOE (green). (**d**) mRNA levels of astrocyte markers: *S100β*, *GFAP*, *VIM*, *AQU4*, *ACSBG1*, and *APOE* in hiPSC-astrocytes and pAstrocytes. Primer sequences are listed in **Supplementary Table 3**. (**e**) mRNA levels of neuronal markers: *TUJ1*, *MAP2AB*, *RELN* and *CACNA1C* in hiPSC-astrocytes and pAstrocytes, relative to h*NGN2*-induced neurons. Primer sequences are listed in **Supplementary Table 3**. (**f**) Principal component analysis of lineage specific marker expression in 23 pairs of hiPSC-astrocytes and isogenic source NPCs, together with four pAstrocyte lines isolated from fetal cerebral cortex, midbrain, cerebellum and whole brain. (**g**) Flow cytometry analysis of S100β (top) and GFAP (bottom) for pAstrocytes from the cerebral cortex, midbrain, cerebellum and whole brain. CC: cerebral cortex, Mid: Midbrain, Cereb: Cerebellum, Mixed: whole brain. hiPSC-Astros: hiPSC-astrocytes, pAstros: primary human fetal astrocytes. Data are represented as mean ± standard deviation. A two-tailed homoscedastic Student’s t-test, n.s.: not significant, * p < 0.05, ** p < 0.01, *** p < 0.001.

We next tested the efficacy of this protocol across 42 NPC lines from 30 individuals (16 male and 14 female) generated as part of three unique hiPSC cohorts (reprogrammed and differentiated through different protocols in independent laboratories) (**Table 2**; **Supplementary Table 2**). All 42 NPC lines ultimately yielded hiPSC-astrocytes (although not necessarily on the first attempt), with an average composition of 90% S100β+ and 82% GFAP+ cells by flow cytometry (**Fig.1b**), and confirmed by immunocytochemistry, although GFAP intensity was highly variable (**Fig. 1a,c**; **Supplementary Fig.1j-k**). Within 30 days, hiPSC-astrocytes were also immuno-positive for the astrocyte markers ALDH1L1^34^ and Vimentin (VIM)^35^, as well as the glutamate transporters GLAST (EAAT1) ^36^ (**Fig. 1c**; **Supplementary Fig. 1l-m**). Compared to NPCs or excitatory neurons (h*NGN2*-induced neurons)^37^, hiPSC-astrocytes expressed high levels of *GFAP*^33^, *S100β*^33^, *VIM*^35^, *AQU4*^38^, *ACSBG1*^25^ and *APOE*^39^ by qPCR (**Fig. 1d**). In addition, hiPSC-astrocytes expressed low levels of the neuronal markers *TUJ1*, *MAP2AB*, *RELN*, and *CACNA1C* (**Fig. 1e**).

**Table 2.** Summary of number of cell lines tested by each assay.

| Cell types |  | Primary astrocytes | hiPSC-astrocytes | NPCs |
| --- | --- | --- | --- | --- |
| Total number of tested lines |  | 4 | 42 | 31 |
| Number of lines tested for each assay | IF | 4 | 13 | 13 |
|  | qRT-PCR | 4 | 29 | 31 |
|  | FACS | 4 | 40 | 13 |
|  | IL-6 ELISA | 4 | 13 | 2 |
|  | RNAseq | 2 | 4 | 6 |

Across a larger panel of neural lineage markers (**Supplementary Table 4**) in 23 hiPSC-astrocyte lines and their isogenic NPC lines, principal component analysis (PCA) revealed that the NPCs grouped together, while the astrocytes (both hiPSC-derived and primary human fetal astrocytes) were more dispersed (**Fig. 1f**; **Supplementary Fig. 1n**). Because it was apparent in our hiPSC-astrocytes as well as the primary human fetal astrocytes (obtained from different donors and brain regions), this variability in lineage marker expression may reflect inter-individual variability and/or differences in regional patterning (**Fig. 1g**; **Supplementary Fig. 1o**).

### hiPSC-astrocytes and primary human fetal astrocytes share similar transcriptional profiles

To query the extent to which global gene expression in hiPSC-astrocytes resembles primary human fetal brain astrocytes, we performed transcriptomic analyses (RNAseq) on hiPSC-astrocytes and primary human fetal astrocytes from two brain regions (cerebral cortex and midbrain), together with isogenic hiPSC-derived NPCs and neurons (**Fig. 2; Supplementary Fig. 2**), comparing them to *in vivo* human and rodent astrocyte transcriptomic datasets (**Figs. 2-3**; **Supplementary Figs. 2-3**). hiPSC-astrocytes showed transcriptional profiles most similar to fetal brain astrocytes. Using principal component analysis and hierarchical clustering, all four hiPSC-astrocytes clustered together with the primary human fetal astrocytes and distinct from the NPC and neuron clusters (**Fig. 2a-b**; **Supplementary Fig. 2a**).

**Figure 2.**
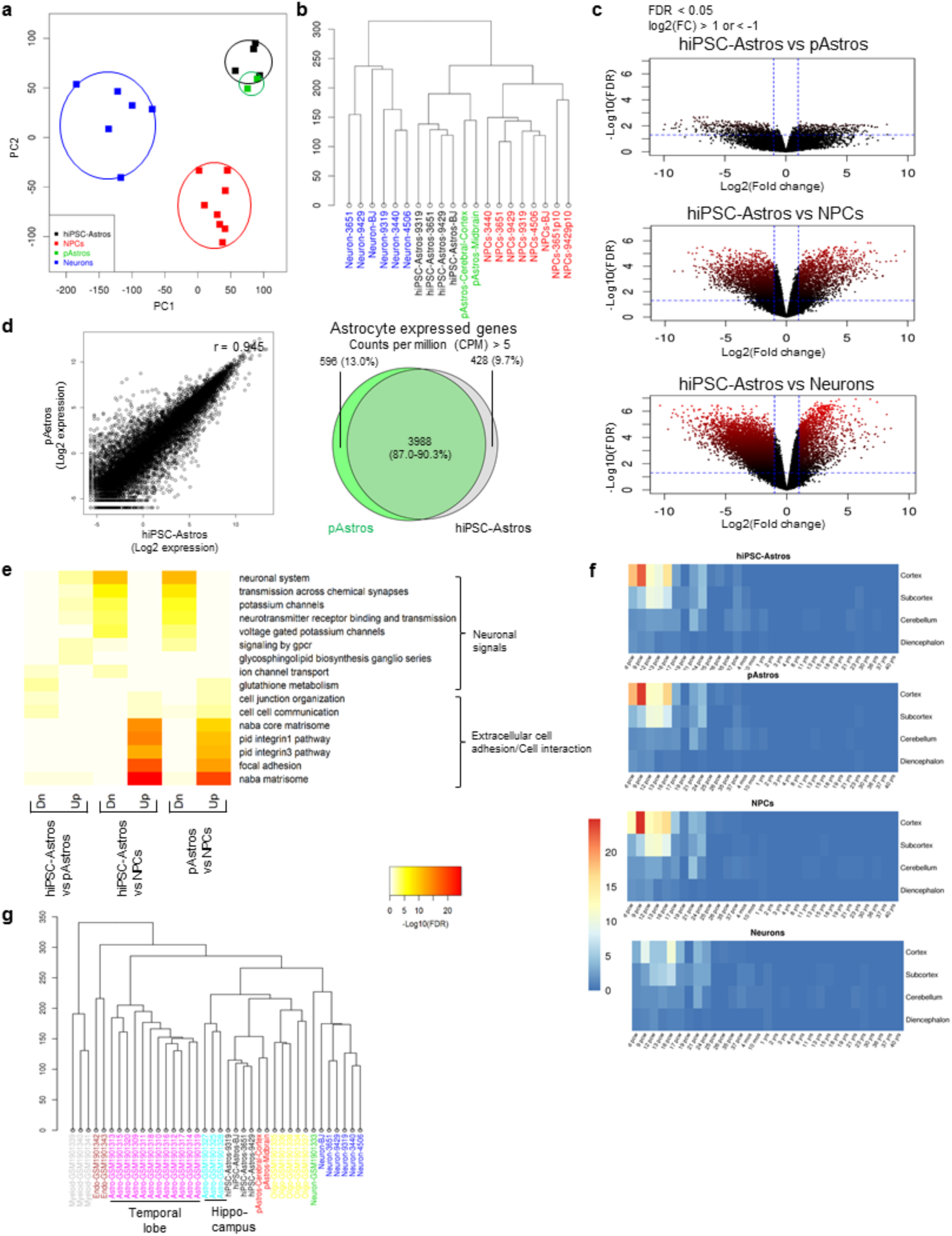
Transcriptional profile of hiPSC-astrocytes and primary human fetal astrocytes. RNAseq analysis of hiPSC-derived NPCs (N=8), neurons (N=6) and astrocytes (N=4) together with pAstrocytes from human fetal cerebral cortex and midbrain. (**a-b**) Principal component analysis (**a**) and clustering diagram (**b**) of hiPSC-derived NPCs, neurons and astrocytes, together with pAstrocytes. (**c**) Volcano plot comparison of hiPSC-astrocytes to pAstrocytes (top), as well as to hiPSC-derived NPCs (middle) and neurons (bottom). Average log2(fold change) versus -log10(FDR) is shown for all genes. Genes upregulated and downregulated by 2-fold change and FDR <0.05 are labeled by red dots. The number of genes differentially expressed between different cell types is indicated by the red color density and quantified in **Supplementary Fig. 2b**. (**d**) Scatter plot (left) comparing gene expression in hiPSC-astrocytes and pAstrocytes. r represents the Spearman correlation coefficient. Venn diagram (right) of overlapping gene expression (CPM>5) between hiPSC-astrocytes and pAstrocytes. (**e**) Functional pathway enrichment analysis of differentially expressed genes between hiPSC-astrocytes and pAstrocytes (left), hiPSC-astrocytes (middle) and NPCs, and pAstrocytes and NPCs (right); hiPSC-astrocytes and pAstrocytes express increased extracellular cell communication signals, but decreased neuronal signals relative to NPCs. (**f**) Heatmap produced by Wilcoxon’s rank-sum comparisons of hiPSC-derived NPCs, neurons and astrocytes, as well as pAstrocytes, relative to the Allen BrainSpan Atlas. (**g**) Fold enrichment from functional pathway analysis of astrocyte enriched genes, group G2 sorted from top 100 most variable genes from **Supplementary Fig. 3a** and **Table 3**.

There were nearly six-fold fewer differentially expressed genes (DEGs) between hiPSC-astrocytes and primary human fetal astrocytes (900 genes) than between hiPSC-astrocytes and isogenic hiPSC-derived neurons (10,000 genes) or NPCs (5500 genes) (**Fig. 2c; Supplementary Fig. 2b**). hiPSC-astrocytes are highly similar to primary human fetal astrocytes (r = 0.945); the majority of both expressed genes (counts per million (CPM) > 1) and enriched genes (counts per million (CPM) > 5) were shared between hiPSC-astrocytes and primary human fetal astrocytes (87.0 and 90.3%, respectively) (**Fig. 2d**). Functional enrichment analyses (using MSigDB) demonstrated that signals regulating neuronal maturation, such as synapse or ion channel formation, were downregulated in hiPSC-astrocytes and primary human fetal astrocytes, whereas signals promoting extracellular cell adhesion and interaction were upregulated (**Fig. 2e**). When we specifically considered just the top 100 most variable genes distinguishing hiPSC-derived astrocytes from NPCs and neurons, functional enrichment analysis identified a group of 19 genes related to reactivity, cytokine, interferon, T cell receptor (TCR), and antigen processing signaling that were enriched in astrocytes (G2) (**Fig. 3a**; **Table 3**; **Supplementary Fig. 3a**).

**Figure 3.**
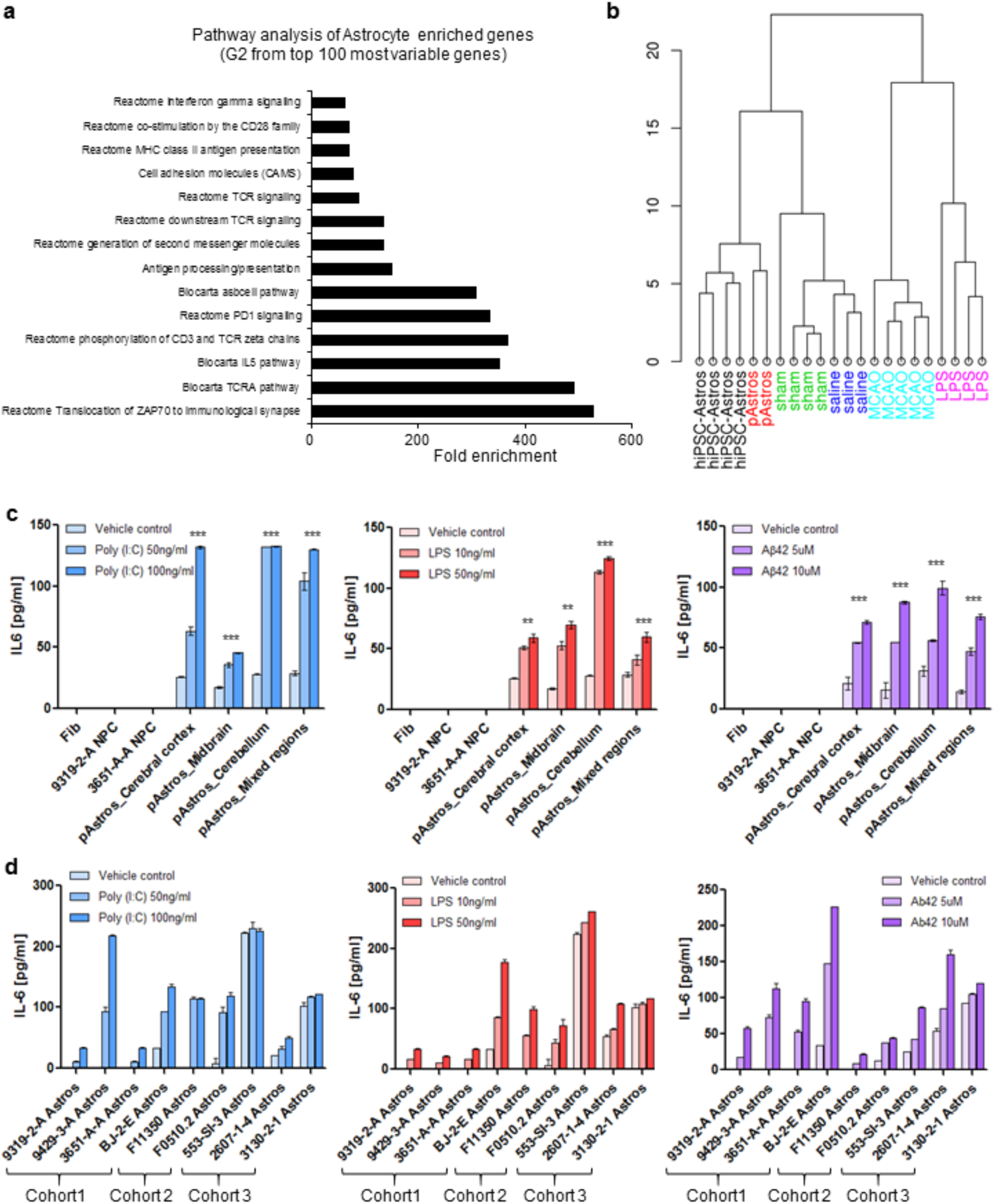
Characterization of the neuroinflammatory status and reactivity of hiPSC-astrocytes. (**a**) Cluster analysis of hiPSC-derived NPCs, neurons and astrocytes, as well as pAstrocytes, combined with a human adult brain data set^41^. hiPSC-Astros: hiPSC-astrocytes, pAstros: primary human fetal astrocytes, FDR: false discovery rate, CPM: counts per million. (**b**) Cluster diagram of hiPSC-astrocytes and pAstrocytes compared to the astrocyte reactivity dataset^42^, which was sorted by reactivity genes enriched in the A1 (LPS-treatment), A2 (MCAO ischemia) and pan-reactive phenotypes, or related controls (saline and sham, respectively). (**c**) IL-6 secretion from pAstrocytes (cerebral cortex, midbrain, cerebellum and whole brain (mixed regions)), and negative controls (fibroblasts and NPCs), after 24-hour treatment with 50 ng/ml or 100 ng/ml of Poly(I:C), 10 ng/ml or 50ng/ml of LPS, or 5 μM or 10 μM of Aβ42 and vehicle controls (saline for Poly(I:C) and LPS and Tris-HCl (pH 8) for Aβ(1-42)), as measured by ELISA. (**d**) IL-6 secretion following 24-hour treatment with Poly(I:C), LPS and Aβ42 across hiPSC-astrocyte differentiations from 9 NPC lines from three independent hiPSC cohorts. hiPSC-Astros: hiPSC-astrocytes, pAstros: primary human fetal astrocytes. Data are represented as mean ± standard deviation. One-way ANOVA with Tukey multiple comparison test, * p < 0.05, ** p < 0.01, *** p < 0.001.

**Table 3.** Annotation of the top 100 most variable genes.

| MSigDB | Group | Overlap size | Group size | MSigDB size | FE | P-value | FDR | Overlap Genes |
| --- | --- | --- | --- | --- | --- | --- | --- | --- |
| Antigen processing and presentation | G2 | 7 | 19 | 57 | 151.3 | 1.71E-14 | 8.49E-11 | CD74;HLA-DOA;HLA-DQA1;HLA-DQB1;HLA-DRA;HLA-DRB1;HLA-DRB5 |
| Cell adhesion molecules (CAMS) | G2 | 6 | 19 | 115 | 64.3 | 3.17E-10 | 3.61E-07 | HLA-DOA;HLA-DQA1;HLA-DQB1;HLA-DRA;HLA-DRB1;HLA-DRB5 |
| Reactome MHC class II antigen presentation | G2 | 5 | 19 | 86 | 71.6 | 6.64E-09 | 6.94E-06 | CD74;HLA-DOA;HLA-DQA1;HLA-DRB1;HLA-DRB5 |
| Reactome translocation of ZAP70 to immunological synapse | G2 | 3 | 19 | 7 | 528 | 1.58E-08 | 1.53E-05 | HLA-DQA1;HLA-DRB1;HLA-DRB5 |
| Reactome phosphorylation of CD3 and TCR zeta chains | G2 | 3 | 19 | 10 | 369.6 | 5.42E-08 | 4.85E-05 | HLA-DQA1;HLA-DRB1;HLA-DRB5 |
| Reactome PD1 signaling | G2 | 3 | 19 | 11 | 336 | 7.45E-08 | 6.22E-05 | HLA-DQA1;HLA-DRB1;HLA-DRB5 |
| Reactome generation of second messenger molecules | G2 | 3 | 19 | 18 | 205.3 | 3.67E-07 | 0.000287 | HLA-DQA1;HLA-DRB1;HLA-DRB5 |
| Reactome downstream TCR signaling | G2 | 3 | 19 | 27 | 136.9 | 1.31E-06 | 0.000965 | HLA-DQA1;HLA-DRB1;HLA-DRB5 |
| Neuron differentiation | G1 | 4 | 28 | 73 | 45.8 | 1.68E-06 | 0.001205 | CNTN4;LMX1B;OTX2;SLIT1 |
| Generation of neurons | G1 | 4 | 28 | 79 | 42.3 | 2.31E-06 | 0.001574 | CNTN4;LMX1B;OTX2;SLIT1 |
| Axon guidance | G1 | 3 | 28 | 22 | 114 | 2.32E-06 | 0.001574 | CNTN4;OTX2;SLIT1 |
| Neurogenesis | G1 | 4 | 28 | 89 | 37.6 | 3.73E-06 | 0.002458 | CNTN4;LMX1B;OTX2;SLIT1 |
| Nervous system development | G1 | 6 | 28 | 366 | 13.7 | 3.95E-06 | 0.002538 | CNTN4;LMX1B;LY6H;OTX2;SHOX2;SLIT1 |
| Reactome TCR signaling | G2 | 3 | 19 | 41 | 90.1 | 4.74E-06 | 0.002876 | HLA-DQA1;HLA-DRB1;HLA-DRB5 |
| Nervous system development | S1 | 5 | 17 | 366 | 18.8 | 4.82E-06 | 0.002876 | CNTN4;LMX1B;LY6H;SHOX2;SLIT1 |
| Biocarta TCRA pathway | G2 | 2 | 19 | 5 | 492.8 | 6.23E-06 | 0.003549 | HLA-DRA;HLA-DRB1 |
| Reactome co-stimulation by the CD28 family | G2 | 3 | 19 | 52 | 71.1 | 9.77E-06 | 0.005322 | HLA-DQA1;HLA-DRB1;HLA-DRB5 |
| Reactome interferon gamma signaling | G2 | 3 | 19 | 57 | 64.8 | 1.29E-05 | 0.006553 | HLA-DQA1;HLA-DRB1;HLA-DRB5 |
| Biocarta IL5 pathway | G2 | 2 | 19 | 7 | 352 | 1.31E-05 | 0.006553 | HLA-DRA;HLA-DRB1 |
| Axonogenesis | G1 | 3 | 28 | 42 | 59.7 | 1.71E-05 | 0.007939 | CNTN4;OTX2;SLIT1 |
| Biocarta asbcell pathway | G2 | 2 | 19 | 8 | 308 | 1.74E-05 | 0.007939 | HLA-DRA;HLA-DRB1 |
| Neuron differentiation | S1 | 3 | 17 | 73 | 56.6 | 1.92E-05 | 0.00858 | CNTN4;LMX1B;SLIT1 |

MSigDB (http://software.broadinstitute.org/gsea/msigdb), is a collection of gene annotations, including gene ontology, and functional pathways. Overlap size, the number of genes shared between a query gene set (eg, genes in group 1, G1) and a gene set in MSigDB (eg, antigen processing and presentation). Group size, the number of genes in a query gene set. MSigDB size, the number of genes in MSigDB gene set. FE: fold enrichment, FDR: false discovery rate.

To assess the extent to which hiPSC-astrocytes are closely related to human brain astrocytes *in vivo*, we compared expression profiles of hiPSC-derived astrocytes, NPCs, neurons, and primary human fetal astrocytes to the Allen BrainSpan: Atlas of the Developing Human Brain (http://www.brainspan.org)^40^ using a Spearman Rank Correlation analysis. Heatmaps generated through a Wilcoxon’s rank-sum test revealed that gene expression in hiPSC-astrocytes and primary human fetal astrocytes was highly correlated with gene expression in human fetal cortical brain tissue (8-16 weeks post conception) (**Fig. 2f**). Unsupervised hierarchical cell type specific cluster analysis demonstrated that hiPSC-astrocytes closely cluster with astrocytes purified by immunopanning from human brain tissue, particularly hippocampal astrocytes, rather than oligodendrocytes, endothelial cells, myeloid cells or neurons^41^ (**Fig. 3a**); pathway analysis further revealed that many astrocyte-enriched genes related to immune signaling (**Fig. 2g**).

### Expression profile of hiPSC-astrocytes resembles quiescent astrocytes

To determine whether the gene expression pattern of our hiPSC-astrocytes more resembles quiescent or reactive astrocytes, we compared them to an RNAseq dataset of “proinflammatory” A1-type and “immunoregulatory” A2-type murine astrocytes, as well as their saline- and sham-treated controls^42^. (Because these are *in vivo* cell types, whereby A1-type astrocytes are induced by a bacterial lipopolysaccharide (LPS) infection and A2-type astrocytes by middle cerebral artery occlusion (MCAO), no comparable dataset exists for human astrocytes.) hiPSC-astrocyte gene expression best clusters with the control conditions (**Fig. 3b**; **Supplementary Fig. 3b**), suggesting that hiPSC-astrocytes may be closer to the quiescent state than to reactive astrocytes.

### hiPSC-astrocytes secrete cytokines in response to inflammatory stimuli

A key effector of the astrocyte neuroinflammatory response in neurodegenerative diseases is interleukin 6 (IL-6)^43^. Consistent with effects observed in primary human fetal astrocytes (**Fig. 3c**), 24-hour treatment with polyinosinic-polycytidylic acid (poly(I:C)) (50 ng/ml or 100 ng/ml), LPS (10 ng/ml or 50 ng/ml), and β-amyloid 42 (Aβ42) (5 μM or 10 μM) led to dose-dependent increases in IL-6 secretion in hiPSC-astrocytes (**Fig. 3d**).

To more fully characterize the release of inflammatory mediators from hiPSC-astrocytes following 24-hour treatment with 5 μM Aβ42, a main component of the amyloid plaques in Alzheimer’s Disease (AD), we measured pro-inflammatory (IL-1β, IL-4, IL-6, IL-8, IL-10, and TNFα) and anti-inflammatory cytokines (IL-1α, IL-2, IL-12, IL-17α, IFNγ, and GM-CSF) by Multi-Analyte ELISArray, confirming that Aβ42 treatment primarily increased IL-6 secretion^44^ (**Supplementary Fig. 3c**), as well as a second pro-inflammatory cytokine, IL-8. We also measured 36 cytokines, chemokines and acute phase proteins using the Proteome Profiler Human Cytokine Array (**Supplementary Table 5**) in baseline conditions and following 24-hour treatment with 5 μM Aβ42 (**Supplementary Fig. 3d-f)**; Aβ42 increased IL-6 release in both hiPSC-astrocytes and primary human fetal astrocytes (**Supplementary Fig. 3d**). Together, our findings indicate that in response to neuroinflammatory cues, hiPSC-astrocytes are capable of secreting pro-inflammatory cytokines.

### hiPSC-astrocytes display phagocytic capacity and promote phagocytic function of microglia

Astrocytes have been reported to phagocytose and degrade β-amyloid in AD^45,46^. We used flow cytometry to examine the ability of hiPSC-astrocytes to phagocytose pHrodo red conjugated myelin or zymosan bioparticles (pHrodo red dye fluoresces red in the acidic environment of phagosomes; zymosan is an agonist of Toll-like receptor 2 (TLR2)^47,48^). hiPSC-astrocytes, primary human fetal astrocytes and BV2 microglia showed a similar capacity to internalize myelin purified from brain homogenates, while zymosan bioparticle uptake was much greater in microglia (consistent with higher levels of TLR2 reported in microglia)^49^ (**Fig. 4a-b**; **Supplementary Fig. 4a-c; Supplementary video 1-3**). The specificity of bioparticle uptake was confirmed by treating cells with cytochalasin D, an inhibitor of β-actin polymerization that reduces phagocytosis (**Fig. 4a-b**).

**Figure 4.**
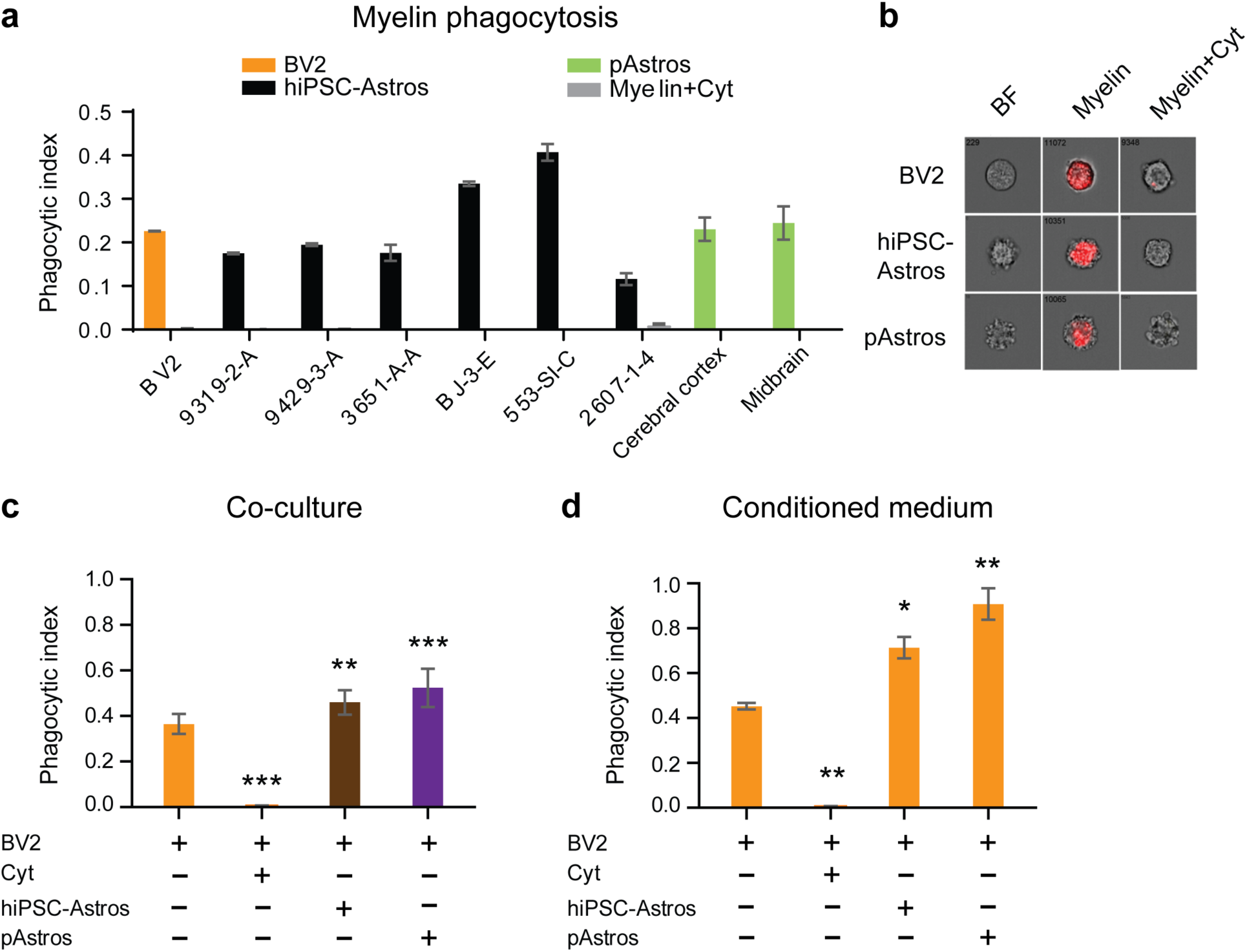
Impact of hiPSC-astrocytes on the phagocytic capacity of BV2 microglial cells. (**a**) Phagocytic indices of BV2 cells, hiPSC-astrocytes, and pAstrocytes incubated with 20 μg of pHrodo-labeled myelin for 3 hours and analyzed by flow cytometry. (**b**) Amnis images representative of pHrodo-red myelin after engulfment in hiPSC-astrocytes, pAstrocytes, and BV2 cells. (**c**) Phagocytic indices of BV2 microglia co-cultured with hiPSC-astrocytes and pAstrocytes that were treated with 30 μg of pHrodo-labeled Zymosan for 3 hours and analyzed by flow cytometry, F(3, 33) = 79.33. (**d**) Phagocytic indices of BV2 microglia treated with astrocyte conditioned medium for 20 hours and then incubated with 30 μg of pHrodo-labeled zymosan for 3 hours for analysis by flow cytometry, F(3, 13) = 30.80. Values in each experiment were averaged from 6 to 8 different control hiPSC-astrocytes and 2 to 4 different pAstrocytes. Data are representative of three independent experiments (n = 3) and shown as mean ± standard deviation. hiPSC-Astros: hiPSC-astrocytes, pAstros: primary human fetal astrocytes. Treatment with 2 μM Cytochalasin D (Cyt) was used as a negative control for phagocytosis inhibition. One-way ANOVA with Dunnett’s post-hoc test, * p < 0.05, ** p < 0.01, *** p < 0.001.

Although microglia are the major phagocytic cells in the brain, astrocytes mediate microglial response to inflammatory stimuli^50-52^. We repeated the pHrodo red zymosan assay using microglia-astrocyte co-cultures; both hiPSC-astrocytes and primary human fetal astrocytes enhanced phagocytic capacity of BV2 mouse microglial cells (**Fig. 4c**; **Supplementary Fig. 4d**). We discriminated microglia and astrocyte phagocytic activity by labeling CD11b- and GFAP-positive cells, respectively (**Fig. 4e**). After 3 hours of labeling, only the CD11b+ BV2 microglial cells phagocytosed the pHrodo red zymosan bioparticles (**Fig. 4e**) Finally, we treated BV2 microglial cells with astrocyte conditioned medium (ACM) for 24-hours prior to analyzing their phagocytic capacity; pre-treatment of BV2 microglial cells with ACM from either hiPSC-astrocytes or primary human fetal astrocytes significantly increased microglial phagocytic capacity (**Fig. 4d**; **Supplementary Fig. 4f**). Taken together, these findings show that both hiPSC-astrocytes and primary human fetal astrocytes secrete factors that increase the capacity of microglia to phagocytose zymosan bioparticles.

### hiPSC-astrocytes display spontaneous calcium transient activity

We next evaluated whether hiPSC-astrocytes exhibit spontaneous calcium transients, as has been described for astrocytes^53^. We used the calcium indicator FLUO-4AM to monitor calcium signaling under basal conditions and in response to a pulse of extracellular glutamate (3 µM)^41^. A single pulse of glutamate produced a slow calcium response in both hiPSC-astrocytes and primary human fetal astrocytes (**Fig. 5**). In addition, some cells exhibited spontaneous calcium spikes, suggesting the presence of a network of connected astrocytes^54^ (**Fig. 5a-d**). To quantify these responses, we measured the frequency of spontaneous activity, the number of spontaneous spikes per time, and the amplitude of the calcium spike (**Fig. 5e-g**). There was no statistical difference in the frequency and number of spikes between hiPSC-astrocytes and primary astrocytes. Interestingly, the amplitude of the spontaneous calcium spike was significantly higher in hiPSC-astrocytes compared to primary astrocytes. Taken together, the excitability of hiPSC-astrocytes was largely indistinguishable from that of primary human astrocytes.

**Figure 5.**
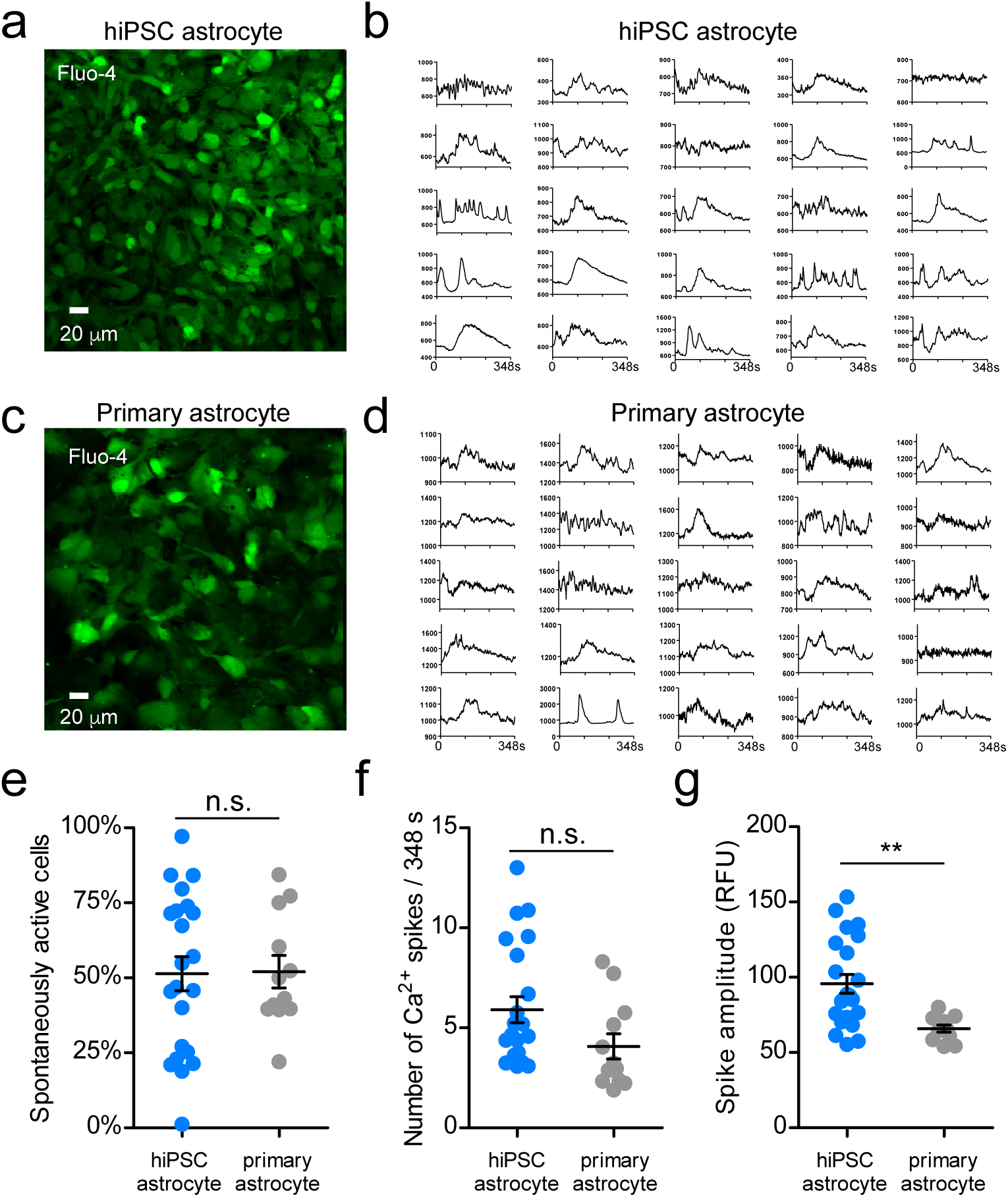
Spontaneous and glutamate-responsive calcium transients in hiPSC-astrocytes and primary astrocytes. (**a,c**) Representative Fluo-4 stained hiPSC-astrocytes and pAstrocytes. Note similarity in shape of cells. (**b,d**) Plots of fluorescence (RFUs) vs. time for 25 ROIs from hiPSC-astrocytes and pAstrocytes. (**e**) Average number of spontaneously active cells per field (318 µm^2^) for hiPSC-astrocytes (N=22 fields from 4 different clones) and pAstrocytes (N=12 fields from 3 different preparations). (**f**) Average number of calcium spikes per 348 s trace for hiPSC-astrocytes (N=22) and pAstrocytes (N=12). (**g**) Average amplitude of calcium spike in each 348s trace for hiPSC-astrocytes (N=22) and pAstrocytes (N=12). Peak excludes the amplitude of the glutamate-induced spike). Each point represents the average multiple ROIs per 318 µm^2^. Line shows mean ± SEM. A two-tailed Student’s t-test, n.s.: not significant, ** p < 0.05. hiPSC-Astros: hiPSC-astrocytes, pAstros: primary human fetal astrocytes.

## DISCUSSION

We screened 11 methods to differentiate hiPSC-derived NPCs into astrocytes, identifying a medium with low FBS (1-2%) that produces homogenous populations of hiPSC-astrocytes within 30-days. Although this is a commercial medium sold for the culture of primary astrocytes, it has not previously been demonstrated to support differentiation of hiPSC to astrocytes. Our method is fast and robust: unlike previous reports, it does not require prolonged culture (∼ 6 months) or a serial sorting process to achieve a pure population of astrocytes^17,22,26^. This protocol was validated across 42 NPC lines from 30 individuals (16 males and 14 females) generated from three independent hiPSC cohorts (**Table 2**; **Supplementary Table 2**). We caution that the quality of the starting NPC population is a critical predictor of success and note that for a few particularly intransient NPC lines, starting from very low passage stocks proved critical to the ultimate successful differentiation of hiPSC-astrocytes. We further note that GFAP seems to be a more variable marker of astrocyte fate (**Fig. 1b**) and recommend instead using S100β when evaluating hiPSC-astrocyte populations. Because our protocols for NPC and astrocyte culture are robust and easily scaled, this methodology is highly amenable to automated culture and future high throughput drug screens using patient-derived astrocytes^55^.

We have established a platform for querying astrocyte-specific contributions to disease predisposition in hiPSC-based models. hiPSC-astrocytes closely resemble primary human fetal astrocytes, purified adult brain astrocytes^41^, and brain tissue homogenate^40^ (**Fig. 2, 3a**). Although the transcriptional profile of our hiPSC-astrocytes most closely resembles a quiescent state (**Fig. 3b**), hiPSC-astrocytes secrete various cytokines and chemokines in response to neuroinflammatory stimuli (**Fig. 3c,d**). hiPSC-astrocytes displayed phagocytic capacity (**Fig. 4a**) and enhanced the phagocytic function of microglia in a co-culture assay (**Fig. 4d**). They also showed spontaneous calcium transient activity and responses to glutamate stimulus (**Fig. 5**). Moreover, because nearly pure populations of excitatory neurons and astrocytes from the same individual, hiPSC clone, and even NPC batch can now be compared, our protocol may reduce experimental variation in studies of the cell autonomous and non-cell autonomous factors underlying disease predisposition.

While many studies of neurodegeneration and synaptic dysfunction use either primary mouse or human fetal glia for human neuron co-culture^56-58^, these current methods are limited by species-specific differences in astrocyte function and limited access to human fetal samples. Our hiPSC-astrocytes overcome these drawbacks, making possible a new hiPSC-based astrocyte-neuron^59^ -microglia^60,61^ co-culture platform for uncovering disease-related mechanisms *in vitro*, which should help to reveal how the cross-talk between these three neural cell types contributes to neurological and psychiatric disease.

## METHODS

Methods and any associated references are available in the online version of the paper.

**Accession Code.** GEO: GSE97904.

*Note: Any Supplementary information and our data are available in the online version of the paper.*

## ACKNOWLEDGEMENTS

BV2 cell line was kindly provided by Marc Diamond (UT Southwestern Medical Center). We thank the continuous support of the Flow Cytometry CORE at the Icahn School of Medicine at Mount Sinai Hospital. The Goate lab is partially funded by the JPB Foundation and the Rainwater Foundation and the Brennand laboratory is partially funded by NIMH R01MH101454, NIA U01P50AG005138-30-1 (Alzheimer’s Disease Research Center: Pilot 30-1), NIH U01AG046170, and the New York Stem Cell Foundation.

## AUTHOR CONTRIBUTIONS

J.TCW, K.J.B. and A.M.G. designed the experiments and wrote the manuscript. J.TCW developed astrocyte differentiation protocol; J.TCW, K.R.B. B.J.H and R.A. differentiated the hiPSC astrocyte lines. M.W. and B.Z. conducted the RNAseq analysis. J.TCW performed the neuroinflammatory assays. A.A.P., J.TCW and S.M. performed the phagocytosis assays. J.TCW, A.A.P. and B.J.H executed flow cytometry analysis on astrocytes and NPCs. J.TCW and K.R.B. performed qRT-PCR and immunofluorescence staining. E.L. and P.A.S. conducted and analyzed the calcium imaging analysis. C.M.K shared prepublication information about astrocyte differentiation condition.

